# Methanotrophy Under Extreme Alkalinity in a Serpentinizing System

**DOI:** 10.1101/2025.09.12.675893

**Authors:** Alta E. G. Howells, Kirt Robinson, Miguel G. Silva, Ellen Cook, Lucas Fifer, Grayson Boyer, Tori Hoehler, Everett L. Shock

## Abstract

Serpentinization produces hyperalkaline, H_2_- and CH_4_-rich fluids that support microbial life in extreme conditions and serve as analogs for ocean worlds such as Enceladus. While methane production in these systems has been well studied, methane *consumption*—especially under high pH—remains poorly understood. Here, we present isotopic, geochemical, and genomic evidence for hyperalkaliphilic (pH > 11) methanotrophy in the Samail Ophiolite in Oman. Using models that account for fluid mixing and gas exsolution, we identify δ^13^CH_4_ enrichment that cannot be explained by abiotic processes alone. The enrichment of ^13^CH_4_ co-occurs with methanotroph 16S rRNA gene sequences, particularly in fluids formed by mixing CH_4_-rich, anoxic fluids with oxidant-rich surface waters. Shotgun metagenomics reveals a metagenome-assembled genome affiliated with *Methylovulum*, encoding a complete methane oxidation pathway, multiple carbon assimilation routes, and Na^+^/H^+^ antiporters—adaptations likely enabling growth above pH 11. Methanotroph diversity and abundance peak in mixed fluids but are suppressed at total ammonia nitrogen concentrations >20 μM. Anaerobic methane-oxidizing archaea (ANME) may also contribute to CH_4_ oxidation in the deep subsurface. Our findings highlight the viability of methanotrophy under extreme alkaline conditions and provide a framework for interpreting δ^13^CH_4_ signals in serpentinizing environments on Earth and beyond.

## Introduction

Serpentinizing systems, where water reacts with ultramafic rocks, are of interest in geoscience as potential resources for geologic H_2_ production and CO_2_ sequestration, in planetary science as analogs for potential targets of solar system exploration, and in microbial ecology as ecosystems to assess novel adaptive strategies to polyextreme conditions. Serpentinization produces H_2_ through the reduction of water coupled to the oxidation of primary ferrous iron minerals to secondary ferric iron minerals^1,2^. Efforts are underway to investigate the potential for geologically produced H_2_ to be mined for fuel^3^. In addition to H_2_ formation, the reacted fluid is hyperalkaline (pH > 11) and rich in calcium. This poises the system to favor carbonate mineral precipitation upon the addition of CO_2_, making serpentinization a natural abiotic process to sequester CO_2_ from the atmosphere^4^. There is evidence for processes like serpentinization occurring on other planetary bodies in our solar system. For example, the Cassini mission detected H_2_ in the plume of Enceladus^5^, which is attributed to hydrothermal water-rock reactions within a likely alkaline subsurface ocean^6^, making serpentinizing systems on Earth a key analog for understanding the potential habitability of Enceladus. The high concentrations of H_2_ that are characteristic of serpentinizing systems could potentially be used as a substrate by a wide range of microorganisms. However, any microbial inhabitants of serpentinizing systems must deal with the stress of hyperalkaline conditions, often in combination with a scarcity of CO_2_ – a unique polyextreme environment with potential to drive novel adaptive strategies.

In addition to high H_2_ concentrations, serpentinizing systems can have high concentrations of CH_4_. The origin of CH_4_ in serpentinizing systems has been attributed to both abiotic processes^7^ and microbial methanogenesis (4H_2_ + CO_2_ → CH_4_ + 2H_2_O)^8^. Both could potentially be stimulated by carbon sequestration (CO_2_ addition) and/or H_2_ production that would serve to produce methanogenic reactants. Resulting increased efflux of the potent greenhouse gas methane, if not captured, could be an unintended adverse consequence of such efforts. In other aquatic systems such as marine waters and sediments, generated methane is often effectively consumed by methane-oxidizing microorganisms (methanotrophs)^9^, raising the question whether a similar consumptive mechanism may occur in serpentinizing systems. Many studies in serpentinizing systems have thus far focused on methane production^8,10,11,12,13,14^. In this study, we addressed the potential for microbial methane consumption.

A combination of geochemical and microbiological measurements was made to assess the occurrence of methanotrophy in the Samail ophiolite in Oman, where uplifted ultramafic mantle rock is undergoing active serpentinization (Leong et al., 2021). This system encompasses numerous wells and natural springs that span a diverse range of fluid chemistry, providing an ideal backdrop against which to assess the conditions that best support methanotrophy. Moreover, existing geochemical, molecular biology, CH_4_ isotope, and microbial activity studies provide extensive chemical and biological context for the present work.

In Oman, subsurface serpentinized fluids sampled through wells and recently drilled boreholes can have δ^13^CH_4_ values as high as 23.9‰^15^. Miller et al. (2018), using methanogen culture enriched from Oman well fluid, showed that such high values can result from methanogenesis under carbon limited conditions^11^. However, Nothaft et al. (2021b) observed the presence of aerobic methanotroph 16S rRNA gene phylotypes in borehole fluids with ^13^C enriched CH_4_, which suggests the enriched δ^13^CH_4_ values could alternatively or additionally be the result of biological methane oxidation^15^. Aerobic methanotrophs have been detected in several studies on serpentinized fluids of Oman. Rempfert et al. (2017)^10^ detected 16S rRNA gene phylotypes of an aerobic methanotroph species, *Methyloccocus capsulatus*, and Kraus et al. (2021) detected RNA transcripts for particulate methane monooxygenase, an enzyme that oxidizes CH_4_ to CH_3_OH in the first step of aerobic methanotrophy^13^. Combined, these results suggest that methanotrophy could contribute to the enriched δ^13^CH_4_ values observed in serpentinizing systems, but previous work has not explored this potential.

In this study, we present an investigation of methanotrophy and its influence on methane carbon isotope fractionation within serpentinizing systems of the Samail ophiolite. To distinguish biotic fractionation from that driven by abiotic processes, we apply isotope fractionation models accounting for physical mechanisms such as fluid mixing and gas exsolution. These models are enabled by the demonstration that total dissolved silicon concentrations serve as a sensitive indicator of the degree of mixing between pristine serpentinized fluids and those in equilibrium with the atmosphere^16,17^. Alongside isotopic modeling, we assess methanotrophic community composition and diversity in sediments associated with serpentinized fluids using shotgun metagenomic and 16S rRNA gene amplicon sequencing. By integrating geochemical modeling with microbial community analysis, we identify the chemical conditions that most strongly support methanotrophic activity in serpentinizing environments.

## Results and Discussion

### Fluid Mixing, A Potential Goldilocks Zone for Methanotrophs

In Oman, reduced, hyperalkaline fluids classified as Type II^16^—enriched in H_2_ and CH_4_— originate from extensive serpentinization in the deep subsurface (>500 meters). These fluids can migrate to the surface through faults and fissures, where they sometimes mix with Type I fluids. Type I fluids have undergone partial serpentinization at shallower depths and are typically circumneutral in pH and contain higher concentrations of electron acceptors such as O_2_ and SO_4_^2− 16^. The mixing of these geochemically distinct fluids generates steep gradients in pH, redox conditions, and substrate availability, creating energetically favorable niches that may support diverse methanotrophic and other chemolithotrophic microbial communities. To illustrate this, we modeled the variation in energetics of methanotrophy that results from conservative mixing between a pristine Type II serpentinized fluid endmember (less than 1% entrainment of Type I fluid, site 140117F) and a Type I oxidized endmember (site 140116B) based on the measured chemical compositions of those fluids (**Supplementary Table 1S)**. With an increasing contribution of Type I serpentinized fluid to the mix, the concentration of methane decreases and the concentrations of O_2_ and SO_4_^-2^ increase (**Fig. 1a**). Relative to either end-member fluid, energy supplies (as kJ per kg fluid) for aerobic methanotrophy (**Fig. 1b**) and anaerobic methanotrophy coupled to SO_4_^-2^ reduction (**Fig. 1c**) are greater in mixed fluids, and peak near the point of stoichiometric equivalence between oxidant and reductant concentrations. These calculations suggest that mixing may be an important physical process for microorganisms that rely on chemical disequilibrium for aerobic and anaerobic methane oxidation. Aerobic methanotrophs in water columns of lakes, ponds, or sediments are known to live at the interface between oxic and anoxic fluids, where they have both a supply of CH_4_ and O_2_^18^. Fluid mixing in Oman provides a similar chemical transition zone and an opportunity for methanotrophs to thrive.

**Figure 1.**
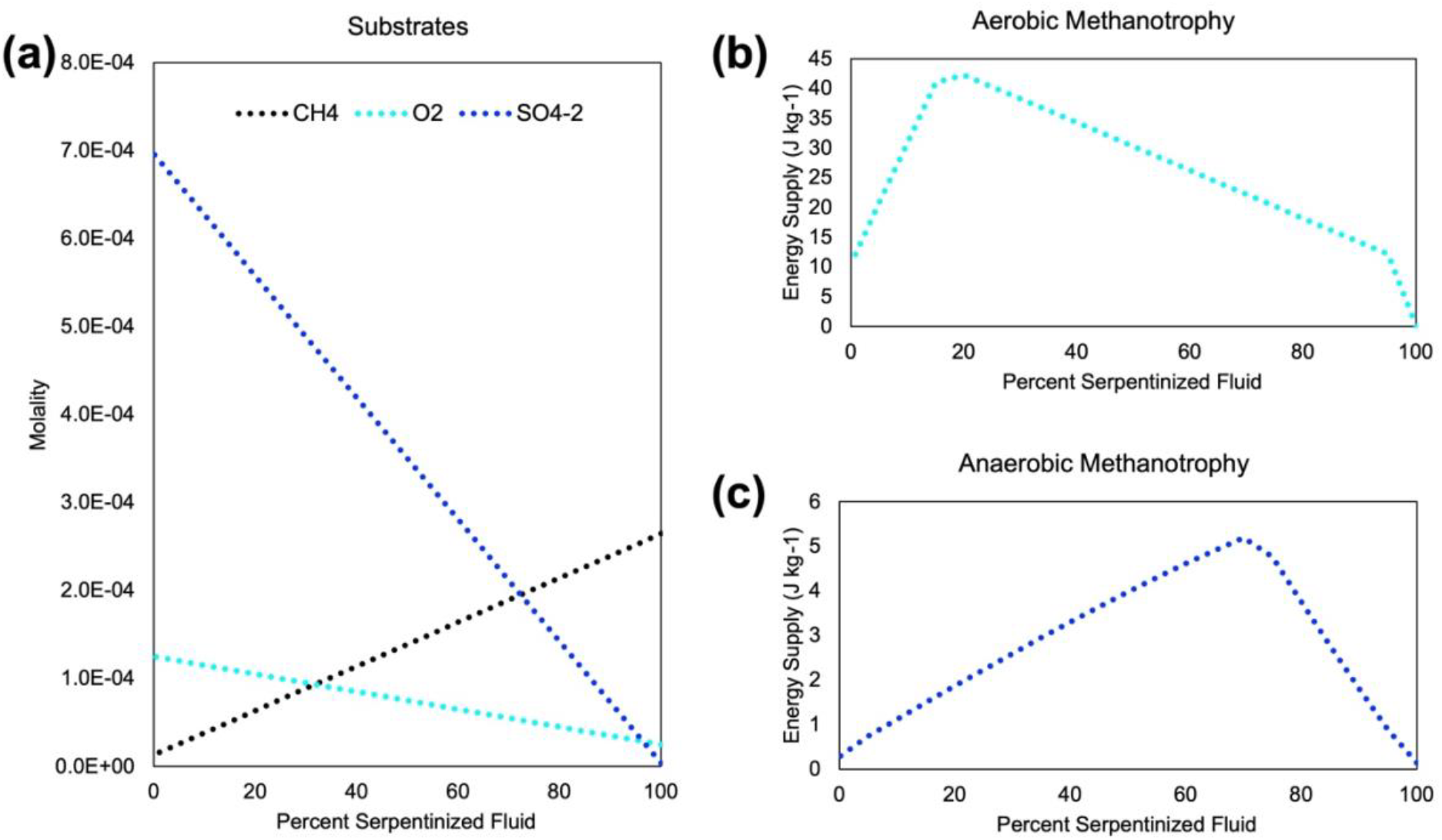
Modeled methane oxidation energetics of fluid mixing. A conservative mixing model was used to estimate the change in fluid chemistry **(a)** and energetics for aerobic **(b)**, CH_4_ + 2O_2_ → CO_2_ + 2H_2_O) and anaerobic methane oxidation **(c)**, SO_4_ ^-2^ is CH_4_ + SO_4_ ^-2^ + 2H^+^ → CO_2_ + H_2_S + 2H_2_O), as two representative endmembers, 140117F for Type II and 140116B for Type I, mix.

### Isotopic Evidence for Methanotroph Activity

Our calculations indicate that methanotrophy would be energetically viable in the systems we evaluated, particularly in zones of fluid mixing. To assess whether methanotrophs are present and active, we compared trends among methane concentrations, methane stable carbon isotopes, and 16S rRNA gene sequences associated with methanotrophs (**Fig. 2**). Preferential consumption of ^12^CH_4_ by methanotrophs results in ^13^C enrichment in the residual (unconsumed) pool of methane, and ^13^C-enriched (“heavy”) methane stable isotope ratios have been used in other studies to infer methanotroph activity. However, other biological and abiotic processes can also influence these ratios. We modeled the influence of fluid mixing and methane exsolution on methane stable isotope ratios and find that they cannot fully account for the level of methane stable isotope enrichment we observed, whereas methanotrophy can. As described below, our observations support active biological methane oxidation in systems with fluid mixing.

**Figure 2.**
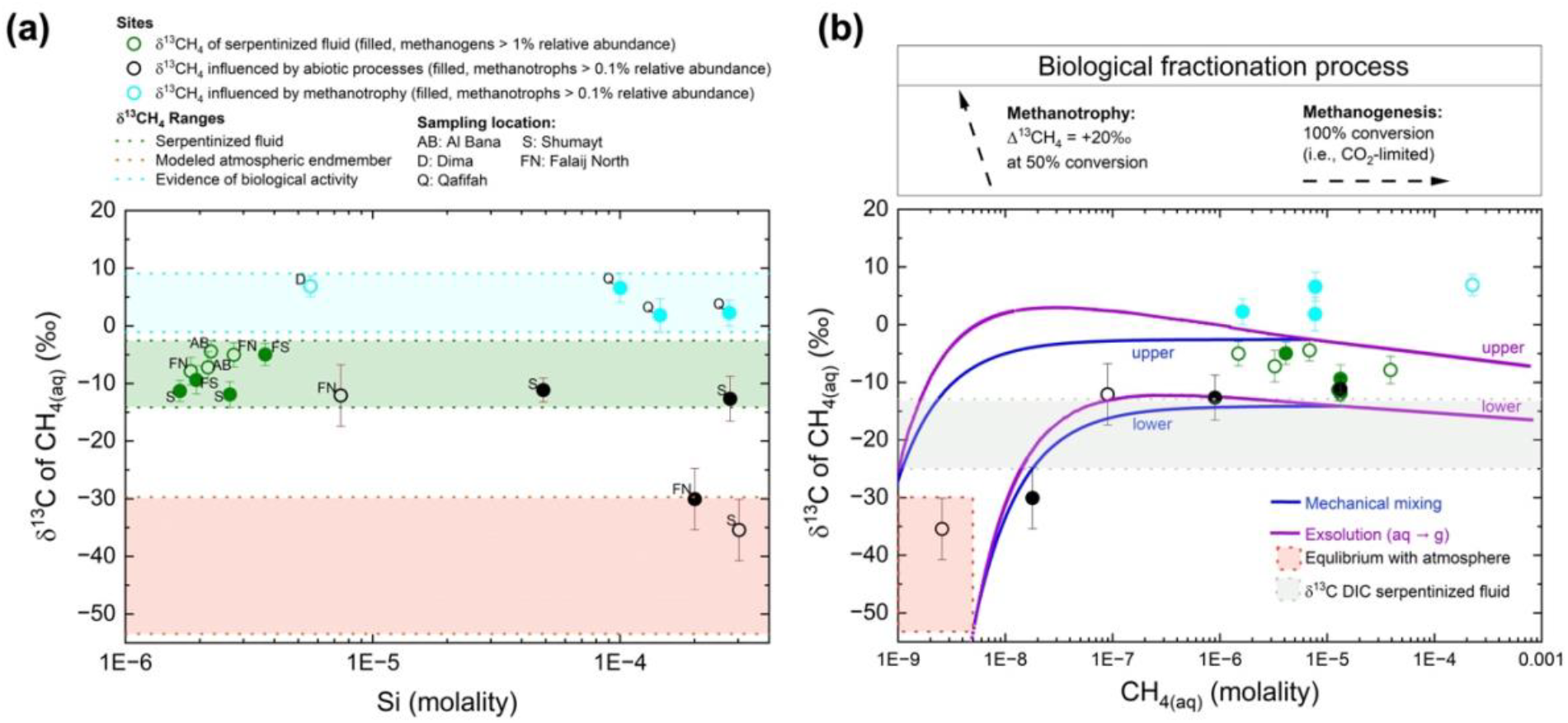
Distinguishing methanotrophic activity from fluid mixing and degassing. δ^13^CH_4_ of dissolved CH_4_ from our sample site fluids is plotted as a function of the concentration of Si in **(a)** and CH_4_ in **(b).** The cyan region in **(a)** depicts the range of δ^13^CH_4_ values whose enrichment in ^13^CH_4_ relative to that of serpentinized fluids (green region) we attribute to methanotrophic activity and is supported by the presence of methanotroph 16S rRNA gene phylotypes (filled in circles). The red region in **(a)** and **(b)** indicates the range of δ^13^CH_4_ measured in the local atmosphere in Oman, which serves as one endmember for the exsolution (purple lines) and mechanical mixing (blue lines) of **(b)**. The other end member for serpentinized fluids is defined by the serpentinized fluids (green region) with the highest **+**1 SD and the lowest **-**1 SD δ^13^CH_4_ values. The grey region in **(b)** is the δ^13^C dissolved inorganic carbon (DIC) range of serpentinized fluids. The box at the top of **(b)** illustrates the direction of fractionation biological activity could impart on the δ^13^CH_4_ values of serpentinized fluid with methanotrophic activity enriching ^13^C^20^ and methanogenesis under carbon-limited conditions imparting no fractionation. Black circles in **(a)** and **(b)** are sites whose δ^13^CH_4_ values are influenced by exsolution and mechanical mixing.

Fractionation models require that we first characterize the CH_4_ carbon isotopic composition of our Type II and Type I fluid endmembers. Near-endmember Type II fluids from four distinct geographic locations (Dima, Qafifah, Shumayt and Falaij North) with <1% of Type I fluid entrained (green circles in **Fig. 2a**) have a narrow range of δ^13^CH_4_ (-12 to -4‰), with the most pristine Type II fluid (lowest Si concentration) having δ^13^CH_4_ = -11.3‰. A sample having the highest Si concentration (little to no contribution of Type II fluid) had a methane concentration of 2.6 nanomolal and methane stable isotope ratio of δ^13^CH_4_ = -35‰, consistent with a fluid in equilibrium with the atmosphere (as calculated from methane concentrations and stable isotope ratios in local air; denoted by orange rectangles in **Fig. 2a** and **2b**).

Our mechanical mixing model is represented by the blue lines in **Fig. 2b**, while purple lines represent the modeled effects of exsolution on methane isotope composition. We base our exsolution model on methods by Knox, 1992 (see *Methods*) which models the transfer of ^12^CH_4_ and ^13^CH_4_ across a liquid/gas interface and predicts enrichments in dissolved ^13^CH_4_ during exsolution^19^. The two lines for each model represent the upper and lower bounds on methane stable isotope ratios that are achievable by that mechanism. Upper bounds are based on application of the models to the highest (most ^13^C-enriched) values of δ^13^CH_4_ observed for Type I and Type II fluid end-members, plus one standard deviation; lower bounds are based on application of the models to the lowest (least ^13^C-enriched) δ^13^CH_4_ measurements of the two endmembers, minus one standard deviation. As shown in **Fig. 2a**, samples that demonstrate intermediate amounts of mixing between Type I and Type II fluids, based on silicon concentration, have wide ranges of δ^13^CH_4_ (-30.1 to 6.9‰) and CH_4_ concentrations (18.0 nanomolal to 226.6 micromolal). As shown in **Fig. 2b**, mechanical mixing generally accounts for combinations of δ^13^CH_4_ and CH_4_ concentration where δ^13^CH_4_ < -8‰, but not fluids with more enriched values.

Four samples are not accounted for by the physical models, all with δ^13^CH_4_ > 1‰. Three out of four fluid samples that are not accounted for by the physical models (**Fig. 2b**) have evidence for abundant methanotrophs (> 0.5% relative abundance), and no methanogens, suggesting their δ^13^CH_4_-enrichments could be the result of preferential consumption of ^12^CH_4_^20^. These sample sites also had visible mixing of Type I fluids into Type II fluids, which provides a source of oxidants (electron acceptors) for methanotrophic metabolisms. Notably, all eight Type II fluid samples that showed less than 1% entrainment of Type I fluids had undetectable or scarce evidence for methanotrophs (< 0.1% relative abundance). We show an example trajectory of δ^13^CH_4_- enrichment during anaerobic microbial methane consumption observed in cultures^20^ (dashed arrow), which could in principle account for the most δ^13^CH_4_-enriched fluids sampled. One sample had a methane stable isotope composition that could not be accounted for by physical processes but had < 0.1% relative abundance of methanotrophs (cyan circle with the lowest concentration of Si in **Fig. 2a**). This sample had no visible signs of mixing at the surface and had the highest measured CH_4_ concentration (**Fig. 2b**), which suggests it had little exposure to the atmosphere, and therefore little potential to entrain atmospheric oxygen. However, previous analysis^16,17^ indicates this sample entrained some Type I fluid (1.46%), perhaps deeper in the subsurface where microbial methane consumption could have caused δ^13^CH_4_-enrichment prior to sampling. Alternatively, as described below, this system may be viable for anaerobic methanotrophs.

### Metagenome Assembled Genome of Methylovolum, a Potential Alkaliphilic Methanotroph

The co-occurrence of methanotroph 16S rRNA gene phylotypes with δ^13^CH_4_ values that cannot be accounted for by physical processes suggests methanotrophy occurs actively in mixed fluids. To assess the functional potential for biological methane oxidation, we conducted shotgun metagenome sequencing on three distinct fluids, all with pH > 11, two of which are Type II fluids with < 1 % influence from mixing, and one with 3.1% entrainment of Type I fluid (96.9 % serpentinized fluid). We used two approaches to resolve metagenome-assembled genomes (MAGs): a co-assemble approach, where sequencing data from all three sites were combined and fed into our pipeline for assembling MAGs, and a “by site” approach, where MAGs were assembled from sequencing data independently for each site. By binning sequencing contigs, we resolved a MAG of an aerobic methanotroph (Bin39) with 96.57% estimated completeness and 0.4% contamination. This methanotroph is classified as belonging to the genus, *Methylovulum*. The co-assembly approach yielded a more complete MAG with more coding sequences than the “by site” approach (Bin28, 95.87% complete), so we focus on the results for Bin39. Using the metabolic map of a *Beijerinkiaceae* methanotroph (MO3_YZ.1)^21^ and the Kyoto Encyclopedia of Genes and Genomes (KEGG)^22^ mapper function as guides, we constructed the core methanotrophic metabolism of our *Methylovulum* MAG (**Fig. 3**). All amino acid sequences of the proteins depicted in **Fig. 3** have 60 % or greater sequence identity to those of previously characterized methanotrophs (see **Supplementary Table 2** for contigs and BLASTp^23^ results). The amino acid sequence for subunit C of pMMO (particulate methane monooxygenase), an enzyme that carries out the first step of methane oxidation – oxidation of methane to methanol – and a phylogenetic marker for methanotrophs, has 94.4 % amino acid sequence identity with *Methylovulum* sp. Within the core metabolic network, we identified a complete pathway for oxidation of methane to formaldehyde, which can feed into the Serine Cycle or RuMP cycle for carbon assimilation, as well as a near-complete H_4_MPT pathway for oxidation of formaldehyde to CO_2_.

**Figure 3.**
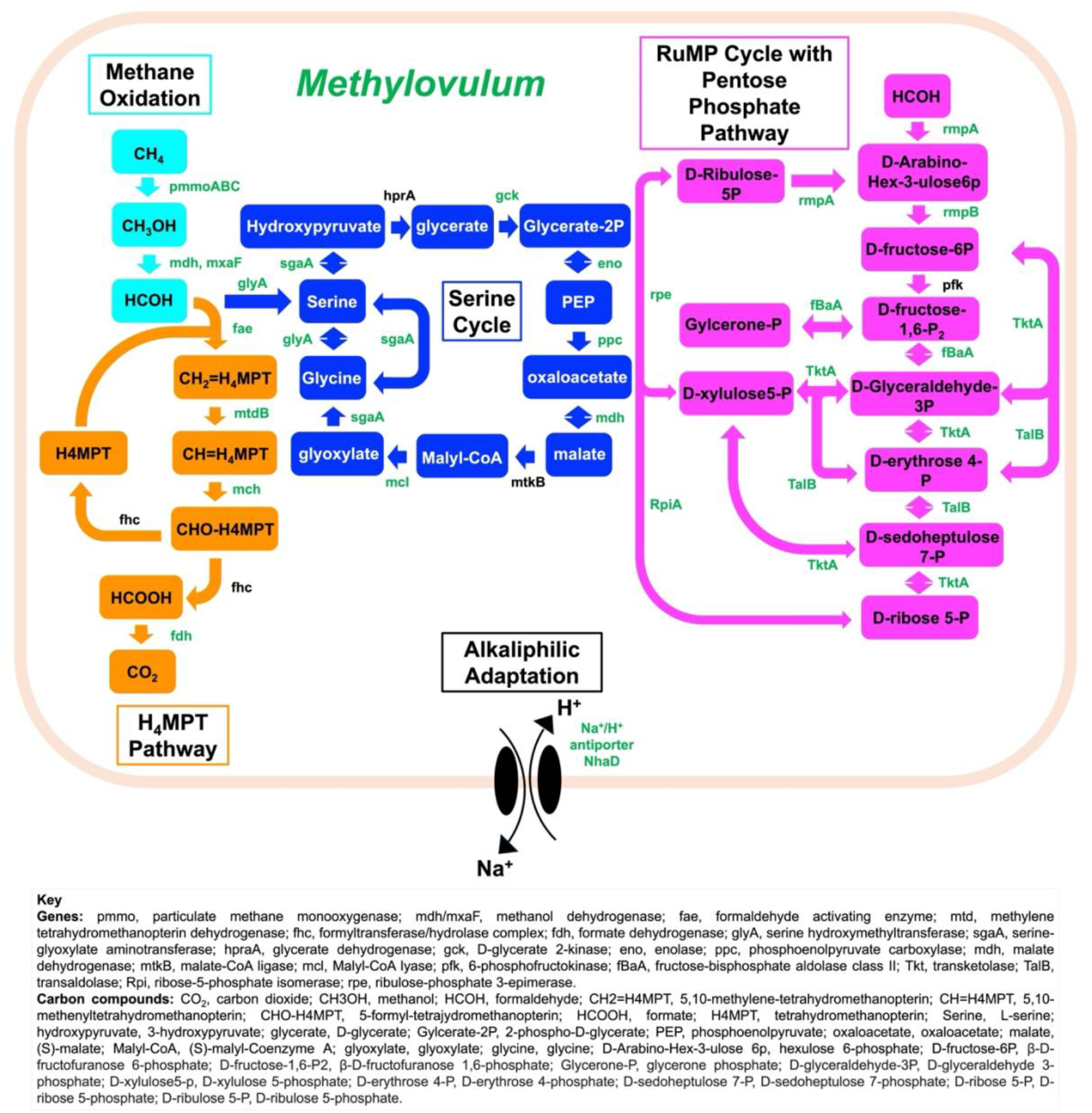
portrays the proposed metabolic pathway of a methanotroph of genus, *Methylovulum*, based on our metagenome assembled genomes, Bin39 and Bin28. Detected enzymes with > 60% sequence identity of cultured methanotroph enzymes are shown in green. All enzymes appear next to the arrow between its substrate and product. Carbon substrates and products appear in colored boxes. Cyan colored arrows and boxes indicate the Methane Oxidation pathway; orange indicates the H4MPT pathway; dark blue indicates the Serine Cycle, and magenta indicates the RuMP Cycle. A gene coding for a Na+/H+ proton antiporter was found, which suggests an adaptation for the potential alkaliphilic species to regulate internal pH in the hyperalkaline serpentinized fluid. Pathways were informed by GhostKOALA and MetaCyc.

The occurrence and abundance of *Methylovolum* corresponds with the enrichment of ^13^CH_4_ (**Fig. 4a** and **4b**). We conducted an analysis to determine the percent of the raw sequencing reads Bin39 composes at each of our metagenome sequencing sites. *Methylovolum* is most abundant at site 140111F (**Fig. 4b**), which has the greatest extent of mixing with Type I fluid among the three shotgun metagenome sequencing sites, and is in a location (Qafifah) with some of the most ^13^CH_4_ ^13^C-enriched δ^13^CH_4_ measurements (**Fig. 4a)**. While δ^13^CH_4_ was not measured for the fluids at site 140111F, it was measured at sites 140111I (67 % serpentinized fluid, δ^13^CH_4_ = 6.6‰) and 140111H (52 % serpentinized fluid, δ^13^CH_4_ = 1.8‰), which are fed by downhill flow of fluid from site 140111F. Given the high relative abundance and diversity of methanotrophs at site 140111F (see **Table 1** and **2** in *Methods*), it is likely that methanotrophic activity by *Methylovulum* contributes to the enriched δ^13^CH_4_ values in the two downstream sites.

**Figure 4.**
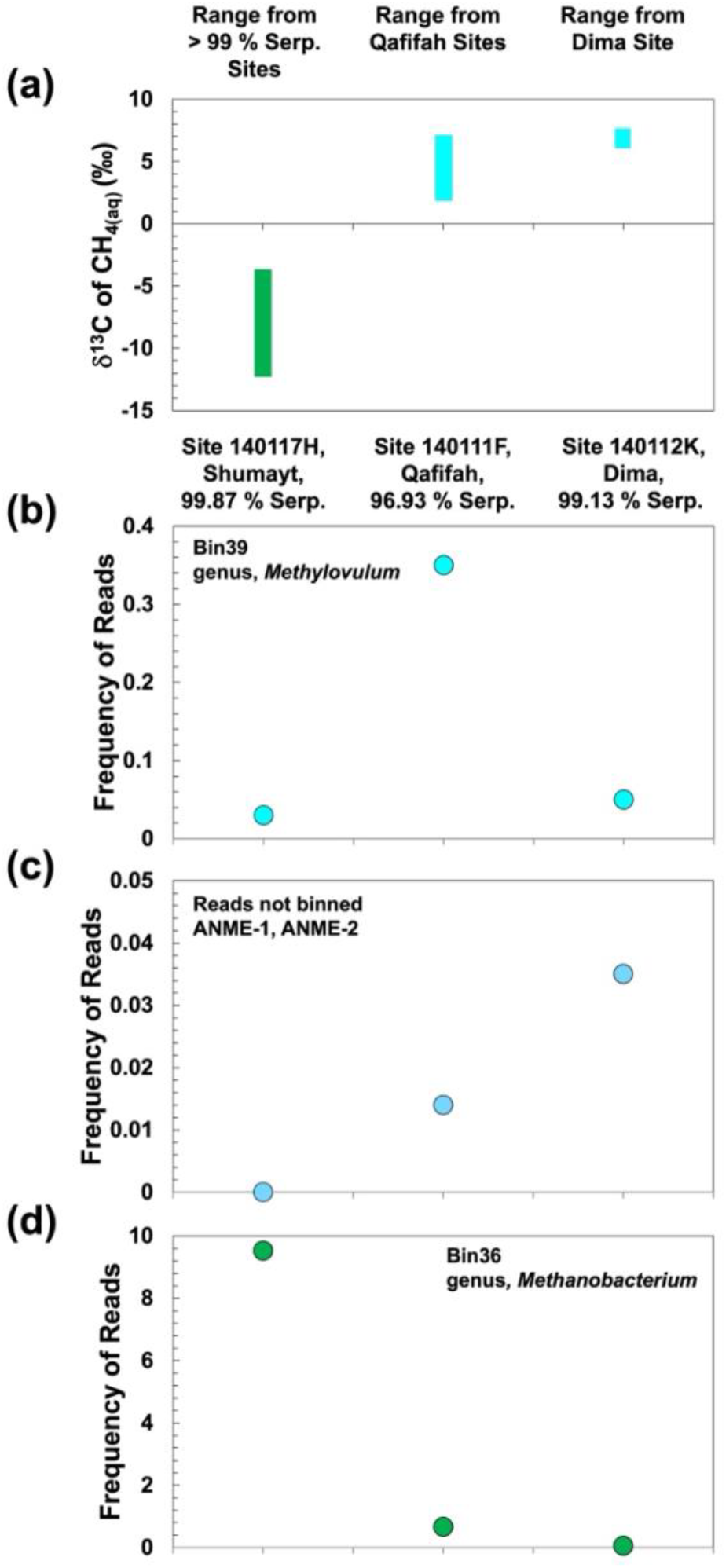
Comparison of d^13^CH_4_ measurements and the frequency of methane cycling taxa based on shotgun metagenome sequencing. **(a)** The range of δ^13^CH_4_ values from sites with > 99% serpentinized fluid composition (green) and the Qafifah (sites 140111G, 140111H and 140111I) and Dima (site 140112M) sites with measurements of δ^13^CH_4_ (cyan). The ranges are defined by + 1 SD of the highest and – 1 SD of the lowest of the δ^13^CH_4_ values from each site group. The relative frequency of mapped reads of the metagenome-assembled genome, Bin39, classified as the aerobic methanotroph genus, *Methylovulum* at a site with > 99 % serpentinized fluid (1401117H), a Qafifah site (140111F) and a Dima site (140112K) is shown in **(b).** The relative frequency of raw metagenome sequencing reads classified as ANME (anaerobic methanotrophs) at each representative site is shown in **(c)**. The relative frequency of mapped reads reported in Howells et al. (2025)^41^ of the metagenome-assembled genome, Bin36, classified as the autotrophic methanogen genus, *Methanobacterium*, is shown in **(d)**.

**Table 1.**
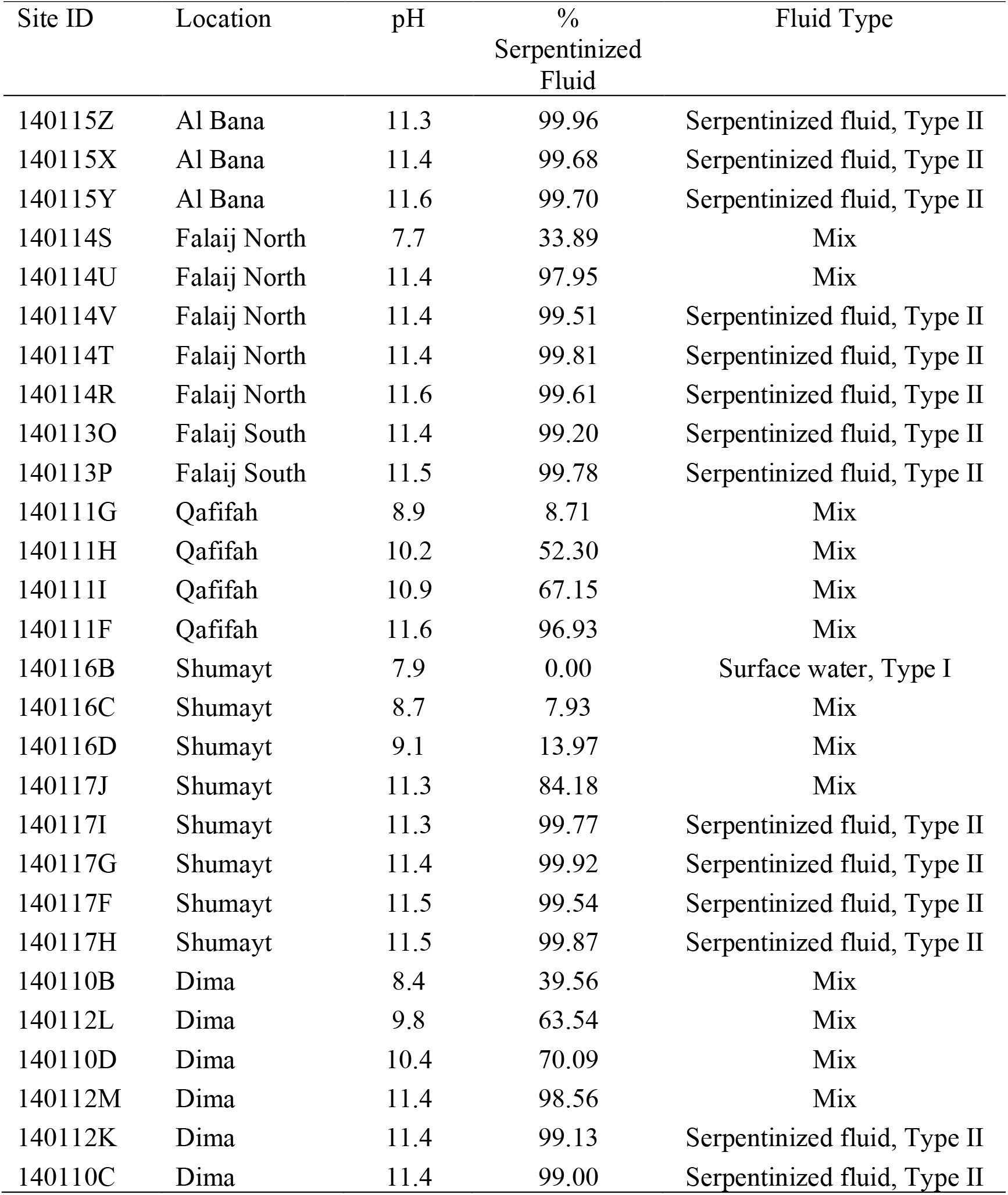
Site description.

**Table 2.** CH_4_ concentration, δ^13^CH_4_ and δ^13^C DIC values, and associated biological assessments.

| Site ID | % Serp.<br>Fluid | CH<sub>4</sub> | δ<sup>13</sup>CH<sub>4</sub> |  | δ<sup>13</sup>C DIC |  | 16S rRNA gene Amplicon<br>Sequencing, |  | Shotgun Metagenome<br>Sequencing |
| --- | --- | --- | --- | --- | --- | --- | --- | --- | --- |
|  |  | nano molality | ‰ | Std.<br>Dev. | ‰ | Std. Dev. | Observed #<br>ASVs | Relative<br>Abundance | Bin ID |
| 140115Z | 99.96 | 18968 | - | - | -17.2 | 0.20 | 3 | 0.1 | - |
| 140115X | 99.68 | 7863 | -4.4 | 1.9 | -17.4 | 0.20 | 0 | < 0.1 | - |
| 140115Y | 99.7 | 3248 | -7.2 | 2.7 | -15.1 | 0.20 | 1 | < 0.1 | - |
| 140114S | 33.89 | 18 | -30.1 | 5.3 | -15 | 0.20 | 9 | 0.1 | - |
| 140114U | 97.95 | 90 | -12.1 | 5.3 | -21.8 | 0.43 | - | - | - |
| 140114V | 99.51 | 1518 | -5 | 2.1 | -24.6 | 0.43 | - | - | - |
| 140114T | 99.81 | 40848 | -7.9 | 2.4 | -16.90 | 0.20 | 1 | < 0.1 | - |
| 140114R | 99.61 | 17227 | - | - | -15.1 | 0.20 | 3 | 0.3 | - |
| 140113O | 99.2 | 4610 | -5 | 1.9 | -20.8 | 0.20 | 1 | < 0.1 | - |
| 140113P | 99.78 | 14378 | -9.4 | 2.4 | -13.1 | 0.20 | 1 | < 0.1 | - |
| 140111G* | 8.71 | 1624 | 2.2 | 2.2 | -14.98 | 0.20 | 14 | 1.3 | - |
| 140111H* | 52.3 | 8003 | 1.8 | 2.9 | -14.0 | 0.20 | 17 | 2.6 | - |
| 140111I* | 67.15 | 7754 | 6.6 | 2.5 | -14.1 | 0.20 | 5 | 0.5 | - |
| 140111F | 96.93 | 157871 | - | - | -9.5 | 0.20 | 1 | 0.1 | Bin39, Bin28 |
| 140116B | 0 | 15 | -35.4 | 5.3 | -14.0 | 0.20 | 3 | 0.1 | - |
| 140116C | 7.93 | 920 | -12.7 | 3.9 | -13 | 0.20 | 4 | 0.4 | - |
| 140116D | 13.97 | 143 | - | - | -13.71 | 0.20 | 2 | 0.2 | - |
| 140117J | 84.18 | 13298 | -11.2 | 2.1 | -14.3 | 0.20 | 2 | 2 | - |
| 140117I | 99.77 | 13612 | - | - | -10.2 | 0.20 | 3 | 0.5 | - |
| 140117G | 99.92 | 9292 | - | - | -18.1 | 0.20 | 1 | 0.2 | - |
| 140117F | 99.54 | 13317 | -11.9 | 2.2 | -15.8 | 0.20 | 1 | < 0.1 | - |
| 140117H | 99.87 | 12728 | -11.3 | 1.9 | -20.2 | 0.20 | 0 | < 0.1 | No Assembly |
| 140110B | 39.56 | 4 | - | - | -9.8 | 0.20 | 2 | 0.1 | - |
| 140112L | 63.54 | 1322 | - | - | -10.2 | 0.20 | 16 | 5.4 | - |
| 140110D | 70.09 | 8912 | - | - | -12.0 | 0.21 | 6 | 2.3 | - |
| 140112M* | 98.56 | 241626 | 6.9 | 1.8 | -4.1 | 0.20 | 1 | < 0.1 | - |
| 140112K | 99.13 | 275897 | - | - | -1.5 | 0.20 | 2 | < 0.1 | No Assembly |
| 140110C | 99 | 204250 | - | - | -6.7 | 0.20 | 0 | < 0.1 | - |

\*Sites where enrichment of <sup>13</sup>CH<sub>4</sub> attributed to biological activity based on fractionation models and biological assessments

Another location in Oman, Dima, has the site with the highest (most ^13^C-enriched) δ^13^CH_4_ among all sites in this study (6.9‰ at 140112M). We detect aerobic methanotrophs at site 140112M in Dima through 16S rRNA gene sequencing, although the relative abundance is < 0.01%. As mentioned above, this location may be suitable for *anaerobic* methanotrophs. We could not resolve a genome of anaerobic methanotrophs, however, raw shotgun metagenome sequence reads were classified through the program Kaiju^24^ and revealed that some reads may be associated with ANME Type I and II anaerobic methanotrophs. The relative frequency of these reads is highest at site 140112K (see **Fig. 4c**), located in Dima. Alternatively, anaerobic methanotrophs could be present and active in the subsurface. In further support of anaerobic methane oxidation by ANME, **Supplementary Fig. 1** shows that sequencing reads classified as ANME have the highest frequency at a site in Dima with the highest energy supply for anaerobic methane oxidation coupled to sulfate reduction, and sampling sites in Dima have the highest occurrence of sulfate reducer 16S rRNA gene phylotypes of the class *Thermodesulfovibrionaceae*, which could serve as partner sulfate reducers for anaerobic methane oxidation. Therefore, the enrichment of ^13^CH_4_ at site 140112M in Dima may be attributed to both aerobic methanotrophs and ANME.

### Methanotroph diversity and distribution

Methanotroph abundance and diversity vary significantly with the extent of fluid mixing. Such mixing can result in variations in pH, oxidant/reductant ratios, and total ammonia nitrogen (TAN) availability, all of which may represent environmental controls on methanotrophs. A broad swath of methanotroph diversity is represented along the mixing gradient, with Verrucomicrobia being more prevalent at sites with < 70% serpentinized fluid and Alphaproteobacteria and Gammaproteobacteria being more prevalent at sites with > 70% serpentinized fluid (**Fig 5**). Also shown in **Fig. 5** is the observed number of ASVs (distinct 16S rRNA gene amplicon sequence variants) classified as aerobic methanotrophs, which can serve as a proxy for the number of unique species of methanotrophs at each site. The observed number of ASVs is lowest in the end member fluids and peaks in mixed fluids, with the highest value being at a site with 52.3% serpentinized fluid. The relative abundance of methanotrophs also peaks in mixed fluids and is highest at a site with 63.5% serpentinized fluid.

**Figure 5.**
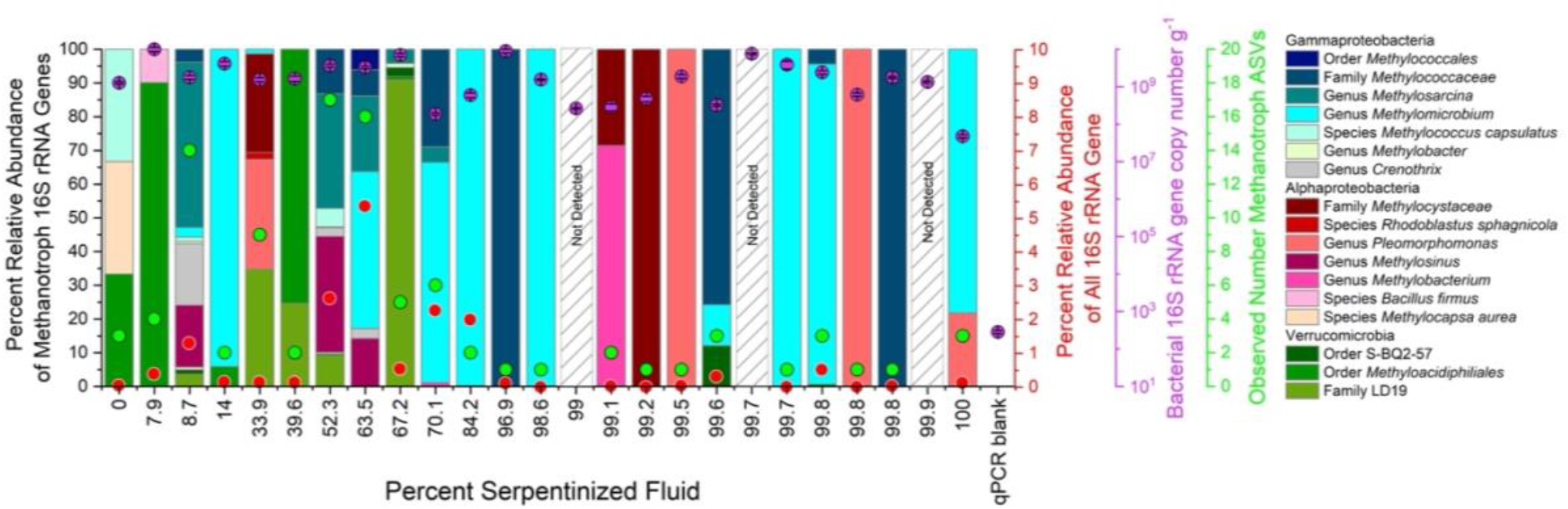
Aerobic methanotroph diversity. The bar-chart depicts the diversity of aerobic methanotroph phylotypes at each site. The relative abundance of methanotroph phylotypes within the whole microbial community is shown by red circles. The relative abundance is approximately equal to abundance given the even copy number of bacteria 16S rRNA genes per gram of sediment (purple circles). The alpha diversity of methanotrophs is shown by the observed number of 16S rRNA gene ASVs classified as aerobic methanotrophs (green circles).

In environmental systems such as lake water columns and peatlands, methanotrophs are commonly found at the oxic/anoxic interface where there is both a supply of O_2_ from the atmosphere and CH_4_ from degradation of organic matter^18,25,26,27,28^. An isolation technique where a CH_4_ and O_2_ counter gradient is established shows that different species of methanotrophs partition across varying levels of CH_4_ and O_2_^29^. We plotted the relative abundance of methanotroph 16S rRNA gene phylotypes with respect to the availability of substrates and energy and find that methanotrophs occupy mixing regions where they have the greatest supply of both CH_4_ and O_2_ and energy (**Fig. 6a**). **Fig. 6a** has been broken down into three zones separated by an energy threshold at 0.7 J kg^-1^ and two reactant ratio thresholds, threshold 1 at log [CH_4_]:[O_2_] ≈ -2.1 and threshold 2 at log [CH_4_]:[O_2_] ≈ 0.4, drawn by the visually observed differences in abundance. On average, the abundance of methanotrophs at sites in Zone 1 is relatively lower (average, ∼0.1%) than sites in Zone 3 (average, ∼0.9%), which have the highest observed abundances, including the maximum abundance (5.4%). The difference in abundance between Zone 1 and Zone 3 corresponds to both a difference in energy supply (low energy, low abundance) and a preference for sites with a relatively higher [CH_4_]:[O_2_] ratio. Zone 1 sites are relatively CH_4_-poor in comparison to sites in Zone 3. Conversely, Zone 2 sites have the highest [CH_4_]:[O_2_] ratios, and yet methanotroph relative abundance is also lower in Zone 2 (average < 0.1%) than in Zone 3. The observed variation in methanotroph abundance and diversity with energy availability and substrate ratio is consistent with that our understanding of methanotroph ecology in other aquatic settings. Here, that variation is driven by mixing of Type I (supply of O_2_) and Type II (supply of CH_4_) fluids that also yield variation in other ecologically important aspects of fluid chemistry.

**Figure 6.**
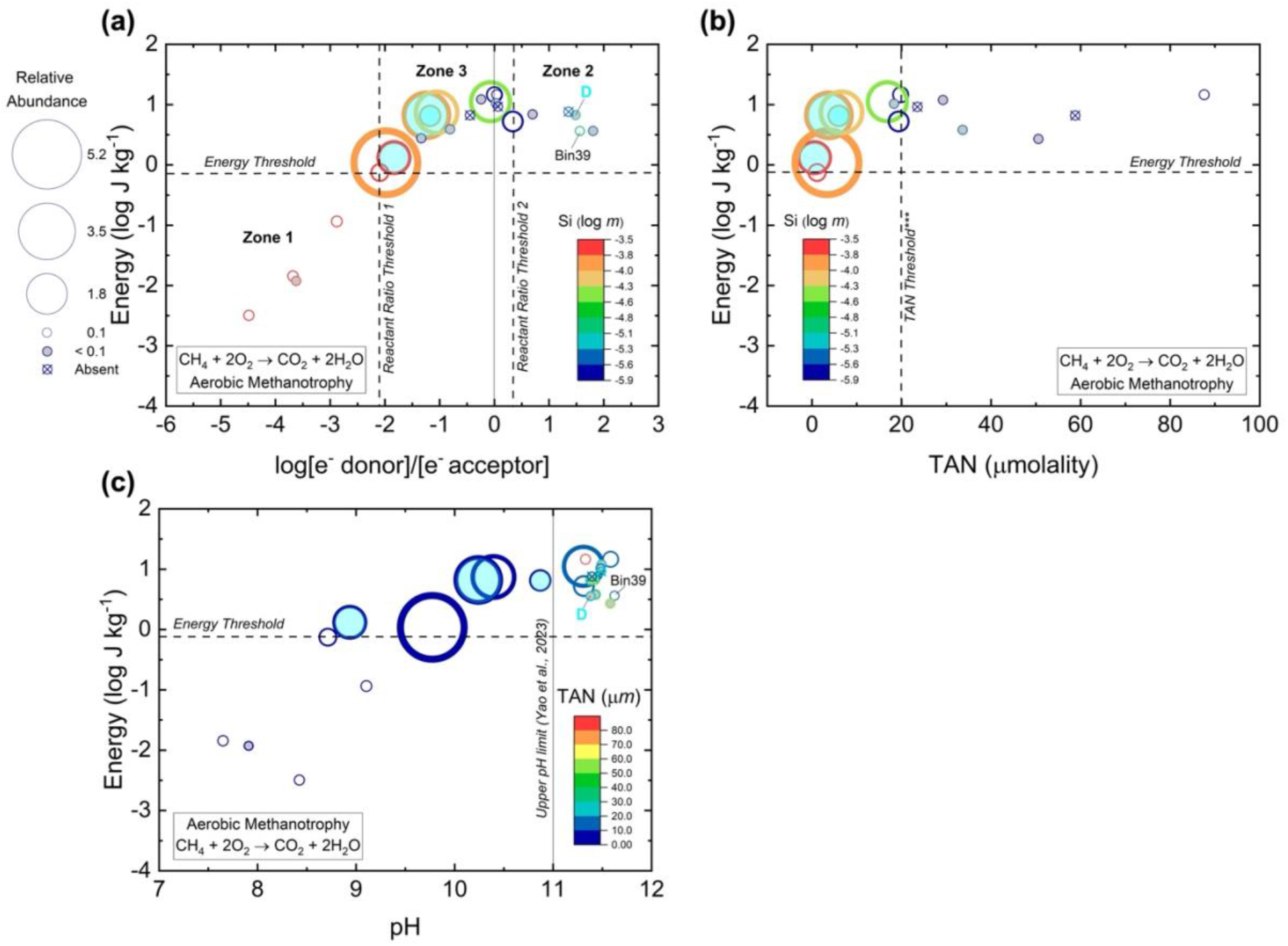
Aerobic methanotroph distribution. The relative abundance of aerobic methanotroph phylotypes is plotted as functions of energy supply and **(a)** the concentration of the electron donor (CH_4_) divided by the concentration of electron acceptor (O_2_) with the concentrations divided by the stoichiometric coefficients of the reactants, **(b)** the concentration of total ammonia nitrogen (TAN) or NH_3_ + NH_4_ and **(c)** pH. For **(a)** and **(b)**, the color scale is the log concentration of Si. For **(c)**, the color scale is the concentration of TAN. An energy threshold was drawn based on observations from this study. The TAN threshold in **(b)** reflects the statistically significant difference in distribution with respect to the concentration of TAN of ∼20 μM. The pH 11 line in **(c)** is the highest pH that aerobic methanotrophs have been cultured in to date^35^. Bubbles filled in with cyan are sites where we observe methanotrophic activity based on the enrichment of ^13^CH_4_ above what can be explained by fractionation due to fluid mixing and exsolution of gas.

Based on previous kinetic studies of methanotroph cultures, NH_3_ are hypothesized to inhibit methanotrophy by competitively binding the active site of particulate methane monooxygenase (pMMO)^30^. In some cases, additions of ammonia nitrogen (reported here as total ammonia nitrogen, NH_3_ + NH_4_^+^, “TAN”) have inhibited methanotroph activity in soils and sediments^31,32,33,34^. While the highest relative abundances of methanotrophs are observed in Zone 3 of **Fig. 6a**, Zone 3 also has some of the lowest abundances, which suggests an additional factor influences methanotroph distribution. TAN concentration could account for this observation. Sites plotted in **Fig 6b** exhibit a dramatic difference in methanotroph abundance above and below a TAN threshold concentration of ∼20 μ*m* (> ∼20 μ*m* average abundance < 0.1%, < ∼20 μ*m* average abundance = 1.4%). The pattern we observe is consistent with inhibition of methanotrophs at higher TAN concentrations, as reported in previous work. To evaluate significance, Mann-Whitney U-tests were conducted. This was done by first making a bar chart of aerobic methanotroph abundance of Zone 3 sites arranged by increasing TAN, **Supplementary Fig. 2**. From there, U-tests were carried out on iterations of group 1 (TAN-poor) and group 2 (TAN-rich), systematically moving group 2 sites into group 1 as the concentration of TAN increases. The descriptive statistics of the abundances for each group (min, max and median) and the U-value and p-value for the tests can be found in **Supplementary Table 3**. Iterative testing was done to be conservative, acknowledging the low sample size, uneven sampling, and error associated with visual observations. The lowest p-value (3.23*10^-4^) is associated with TAN-poor sites that have TAN concentration between 0.57 and 19.83 μ*m*, and TAN-rich sites that have TAN concentrations between 23.53 and 87.60 μ*m*. The lowest p-value suggests that the ∼20 μ*m* threshold accounts for the difference in abundance with the greatest statistical significance. Therefore, aerobic methanotroph activity in serpentinized fluids in Oman are likely inhibited when the concentration of TAN is greater than ∼20 μ*m*.

Solution pH is an additional factor that may influence the growth and reproduction of methanotrophs. The highest pH at which methanotrophs have been cultivated is pH 10.5, and the upper pH limit for methanotrophs is reported as pH 11^35^. In **Fig. 6c**, we assess the distribution as a function of energy and pH with TAN as the color scale. Methanotrophs are distributed across the pH range and are present above pH 11, exceeding the current upper pH limit. The ability of methanotrophs to survive in such hyperalkaline pH regimes may be attributed to possible adaptations observed in this study, such as Na^+^/H^+^ antiporters. For example the detection of *Methylovulum* (Bin39) at a site with pH 11.6 fluid indicates that it is likely an alkaliphile. We found evidence of alkaliphilic adaptations with two genes that code for Na^+^/H^+^ antiporters, type NhaD, which pump Na^+^ out of the cell and H^+^ into the cell^36^. The genes are not identical, with one having 87.36% amino acid sequence identity with an NhaD of *Methylovulum miyakonense* and the other having 72.01% identity with NhaD of *Methylobacter psychrophilus* based on BLASTp results (see **Supplementary Table 2**).

### Connecting to the Subsurface and Beyond

To disentangle the contributions of various processes to δ^13^CH_4_ signatures, we developed a framework that integrates: (1) the isotopic composition of endmember CH_4_, (2) the effects of physical fractionation via fluid mixing and gas exsolution, (3) fractionation associated with biological methane oxidation, and (4) the isotopic composition of dissolved inorganic carbon, which may serve as a substrate for autotrophic methane production. This comprehensive approach allows us to distinguish biological signatures from isotopic patterns typically attributed to abiotic methane sources^7,8,11^.

Our results indicate that the enrichment of ^13^CH_4_ beyond what can be accounted for by physical processes such as fluid mixing and gas exsolution is most likely the result of biological methane oxidation. Methanotrophic activity is closely associated with fluids that exhibit evidence of mixing between Type I fluids (in equilibrium with the atmosphere) and Type II fluids (pristine, deeply sourced serpentinized fluids). This interpretation is supported by the presence of aerobic methanotrophic 16S rRNA gene phylotypes, which are most abundant and diverse in samples from sites with the greatest extent of fluid mixing.

Shotgun metagenomic data further support this conclusion, revealing the presence of the aerobic methanotroph genus *Methylovulum* at a site with some of the highest measured δ^13^CH_4_ values. The metagenome-assembled genome (MAG) of *Methylovulum* encodes a complete methane oxidation pathway, multiple carbon assimilation pathways, and adaptations to hyperalkaline conditions, including two distinct genes encoding Na^+^/H^+^ antiporters.

In addition to aerobic methanotrophs, our data suggest a potential role for anaerobic methane oxidizers (ANME groups I and II). The highest abundance of ANME-affiliated shotgun metagenomic reads was observed at a site with the most enriched δ^13^CH_4_ value, where aerobic methanotrophic 16S rRNA gene phylotypes were nearly absent. These findings suggest that both aerobic and anaerobic methane oxidation may contribute to the isotopic enrichment of CH_4_ in the Samail Ophiolite.

While we do not rule out abiotic processes as contributors to heavy δ^13^CH_4_ values, our findings suggest that, in Oman, these signatures are more plausibly explained by active methanotrophy—particularly in environments where fluid mixing is prevalent and biological evidence is strong.

These insights have broader implications for interpreting δ^13^CH_4_ in other geological settings. Methane isotopic composition has been an important geochemical tool for deciphering the provenance of methane, to include biological overprints, and has been proposed for use in seeking evidence of life elsewhere. Our study highlights the nuance inherent in such an approach, and the need for caution. As we analyze existing δ^13^CH_4_ data from systems such as Mars, and anticipate future measurements at ocean worlds like Enceladus, it is essential to consider physical, chemical, and biological fractionation processes. δ^13^CH_4_ values alone are not definitive biosignatures but can become powerful indicators when interpreted within a multi-line framework that integrates isotopic, geochemical, and microbial evidence.

## Methods

### Sampling Sites

Table 1 describes the sampling sites in this study. The percent serpentinized fluid was calculated using the concentration of Si measured in sampling site fluids and a mixing model as described in Howells et al., 2022. Fluids composed of > 99 % serpentinized fluid are characterized as pristine serpentinized fluid/Type II. The fluid with the highest concentration of Si represents fluid influenced from the surface (surface water/Type I), and sites that have between 0-99% serpentinized fluid are considered mixtures of Type I and Type II fluids. In total, 28 sites from 6 locations (Dima, Qafifah, Shumayt, Falaij North, Falaij South, and Al Bana) are in this study. The aqueous chemistry and dissolved gas concentrations were previously reported in Leong et al. (2021)^16^ and Howells et al. (2022)^17^. For this study we analyzed δ^13^C DIC of fluids at all sites. At 17 sites we analyzed δ^13^CH^4^. Shotgun metagenome sequencing was performed on DNA extracts from sediments of 3 sites. For assessment of methanotroph diversity at each of our study sites we analyzed 16S rRNA gene sequences published in Howells et al. (2022)^17^. For assessment of geochemical parameters that may influence methanotrophs we used geochemical data previously reported in Howells et al. (2022)^17^ and Howells et al. (2023)^37^. The study sites in Howells et al. (2022)^17^ and (2023)^37^ are the same sites, sampled at the same time as this study. An overview of sites with overlapping geochemistry, isotopic analyses, and biological analyses can be found in **Table 2**.

### Isotope Analysis

We analyzed the dissolved gas samples described in Howells et al. (2022)^17^ for the bulk isotopic composition of CH_4_. After analyzing the total concentration of CH_4_ with a gas chromatography flame ionizing detector as described in Howells et al. (2022)^17^, approximately one year later we analyzed the samples for stable carbon isotopes (δ^13^CH_4_). To test for leaks (to avoid unintended fractionation) the samples were first reanalyzed for CH_4_ concentrations and those with significant (> 1 standard deviation) decreases in their concentrations were discarded. Those that had not leaked were injected in triplicate into a Picarro cavity ring-down spectrometer G2201-I equipped with a small-sample isotope module. Ultra Zero Air from Praxair was used as the carrier and for diluting samples and standards; standards were purchased from Air Liquide and 20 mL of each standard was injected in triplicate to produce three-point linear calibration curves with R^2^ values > 0.999, with the following δ^13^CH_4_ values: -69 ± 1, -36 ± 1, and 5 ± 1‰, all at 500 ppmV concentrations. Each sample was diluted to approximately 500 ppmV (when possible) and 20 mL of the resulting diluted sample was injected into the instrument. CH_4_ concentrations and δ^13^CH_4_ values for each site are reported in **Table 2**.

Dissolved inorganic carbon (DIC) measurements were performed using an OI-Analytical total organic carbon (TOC) analyzer coupled to a continuous flow Thermo Electron DeltaPlus Advantage isotope ratio mass spectrometry (IRMS). The setup and methods for the coupled TOC-IRMS were derived from Gilles St-Jean (2003)^38^. Briefly, the method consists of heating, followed by acidification of the sample with phosphoric acid to drive off the DIC as carbon dioxide (CO_2_), which is sent to the IRMS. Data can be found in the **Table 2**.

### Aqueous Chemistry Modeling and Energetic Calculations

For estimating energy supply for aerobic methanotrophy (CH_4_ + 2O_2_ → CO_2_ + 2H_2_O) and anaerobic methanotrophy coupled to sulfate reduction (CH_4_ + SO_4_^-2^ + 2H^+^ → CO_2_ + H_2_S + 2H_2_O) we used the fluid chemistry of sampling sites reported in Howells et al. (2022)^17^ and the Water-Organic-Rock-Microbe (WORM) Portal (https://worm-portal.asu.edu) using the AqEquil Python package^39^ and wrapper for EQ3/6^40^ following methods described in Howells et al. (2025)^41^. **Supplementary Table 1** summarizes the concentrations of reactants and energy supply for each site.

### Isotope Fractionation Modeling

To assess whether ^13^C-enriched CH_4_ in springs could be explained by abiotic physical processes, we applied a *mechanical mixing model* between shallow groundwater (Type I) fluids and serpentinized (Type II) fluids as well as a *methane exsolution model*, whereby ^13^C-enrichment occurs as ^12^CH_4_ is preferentially degassed from CH_4_-rich Type II fluids. For the mechanical mixing model we chose two empirically determined endmember fluids, one being an average of Type II fluids with the lowest dissolved silica that indicated <1% mixing with surface water (see **Table 2**), and the other a calculated Type I fluid saturated with atmosphere based on measurements we made of the local air. To calculate the Type I fluid, we used equilibrium constants for CH_4_ dissolution and δ^13^CH_4_ fractionation obtained from the software SUPCRT92^42^ with supporting thermodynamic data^43,44,45,46^ and a previous study^47^, respectively. With CH_4_ concentration and δ^13^CH_4_ values for both endmember fluids, we simply recalculated those values for different proportions of those fluids for the mechanical mixing model.

For the methane exsolution model, we relied on previous research^19^ and chose parameters that would maximize predicted ^13^C-enrichment in the empirically determined endmember Type II fluid, with the intention of setting an upper limit for abiotic physical processes. See the **Supplementary File** for further details on this calculation.

### 16S rRNA Gene Amplicon Sequencing and Analysis

16S rRNA gene amplicon sequences reported in Howells et al. (2022)^17^ were taxonomically classified using the non-redundant GreenGenes 16S rRNA database^48^. We acknowledge this is an older database, however when comparing taxonomic assignments against those reported in Howells et al. (2023)^37^ which used the Silva database^49^, we found no significant differences. After taxonomic classification of ASVs, aerobic methanotroph phylotypes were identified and further characterized using BLASTn, https://blast.ncbi.nlm.nih.gov/Blast.cgi?PAGE_TYPE=BlastSearch^50^; see **Supplementary Table 4** for those results. After identifying methanotroph ASVs, methanotroph phylotypes were grouped taxonomically, and their relative abundances at each sampling site were calculated (see **Supplementary Tables 5 and 6**).

### Methanotroph Metagenome Assembled Genomes Analysis

For this study we analyzed the metagenome assembled genomes (MAGs) from the shotgun metagenome sequencing analysis described in Howells et al. (2025)^41^. The co-assembly approach described in Howells et al. (2025)^41^ yielded one MAG classified as belonging to the genus, *Methylovolum* (Bin39). The by-site approach yielded one MAG classified as *Methylovolum* from site 140111F (Bin28). Methanotroph genomes could not be resolved from sites 140117H and 140112K. An overview of the assembly statistics is in **Table 3** and the relative abundances of the MAGs were estimated using methods described in Howells et al. (2025)^41^ and are summarized in **Table 4**.

**Table 3.** Assembly statistics for metagenome assemble genomes (MAGs) of methane cycling taxa.

| Bin ID | Method | Site/s | Core Methane Metabolism | Taxonomy (GTDB-Tk) | Est. Compl. | Est. Contam. | GC (%) | Size (bp) | Contig (no.) | CDS |
| --- | --- | --- | --- | --- | --- | --- | --- | --- | --- | --- |
| Bin39 | co | All | Aerobic methanotrophy | Family, *Methylomonadaceae*; Genus, *Methylovulum* | 96.57 | 0.37 | 65 | 2,346,974 | 201 | 2,424 |
| Bin28 | by site | 140111F | Aerobic methanotrophy | Family, *Methylomonadaceae*; Genus, *Methylovulum* | 95.87 | 0.58 | 65.1 | 2,336,462 | 225 | 2,485 |

**Table 4.** Estimated relative abundances of methane cycling taxa based on the mapped reads of MAGs at each site.

| Bin ID | Method | Site/s | Core Methane Metabolism | Taxonomy (GTDB-Tk) | % Mapped Reads (Bowtie) |  |  |
| --- | --- | --- | --- | --- | --- | --- | --- |
|  |  |  |  |  | 140117H, 99.9 % | 140112K, 99.1 % | 140111F, 96.9 % |
| Bin39 | co | All | Aerobic methanotrophy | Family, *Methylomonadaceae*; Genus, *Methylovulum* | 0.03 | 0.05 | 0.35 |
| Bin28 | by site | 140111F | Aerobic methanotrophy | Family, *Methylomonadaceae*; Genus, *Methylovulum* | 0.04 | 0.05 | 0.35 |

### Plotting and Statistical Analyses

OriginPro version 2024^51^ was used to make scatter plots, bubble plots, and bar charts. Given non-normal histogram distributions, we implemented the Mann-Whitney U-test function in OriginPro to assess the statistical significance of distribution differences with respect to geochemical and biological parameters. Given the small sample size of sites, we used the exact p-value with a 0.05 cutoff for significance.

## Data Availability

All original data generated in this study are publicly accessible at the NCBI Sequence Read Archive (SRA) under BioProject ID PRJNA1207711: https://www.ncbi.nlm.nih.gov/sra. Metagenome-assembled genomes (MAGs) produced as part of this work are available from the authors upon reasonable request without restriction.

## Acknowledgments

This research received financial support for its execution, authorship, and publication. Fieldwork was supported by NASA Exobiology Grant NNX12AB38G, the NASA Astrobiology Institute’s Rock-Powered Life (RPL) initiative under Grant NNA15BB02A, and NSF Grant EAR-1515513. This work used the Water-Organic-Rock-Microbe (WORM) Portal funded by NSF Grants EAR-1949030 and EAR-2149016. Shotgun metagenomic sequencing was conducted with funding from the Deep Carbon Observatory’s Census of Deep Life, supported by the Alfred P. Sloan Foundation. Additional support was provided by NASA’s Planetary Science Division through the Center for Life Detection ISFM team. Alta E. G. Howells received postdoctoral fellowship funding through the NASA Postdoctoral Program at NASA Ames Research Center. Lucas Fifer’s contributions were supported by the University of Washington Astrobiology Program. Miguel Silva, and Ellen Cook contributed to the project as part of the BMSIS Young Scientist Program (YSP) at NASA Ames Research Center, under the guidance of Alta E. G. Howells and Tori M. Hoehler.

## Author Contributions

Alta E. G. Howells led the production of this paper, participated in field work and sample collection, carried out 16S rRNA gene sequencing analysis, shotgun metagenome sequencing, dissolved gas concentration analysis, led the intellectual development of this study and led writing. Kirt Robinson participated in field work and sample collection, measured δ^13^C of CH_4_ and dissolved inorganic carbon, developed the bulk isotopic fractionation models of CH_4_, provided intellectual feedback and assisted with writing. Miguel G. Silva annotated and assembled the core metabolisms of the *Methylovolum* MAGs. Ellen Cook helped develop the processing pipeline for shotgun metagenome sequencing data. Lucas Fifer and Grayson Boyer carried out energetic calculations. Tori Hoehler developed energetic calculations for conserved mixing, provided intellectual feedback and assisted with writing. Everett L. Shock was the primary investigator of this research, participated in field work and sample collection, provided intellectual feedback and assisted with writing.

