## Supplementary File for "Methanotrophy Under Extreme Alkalinity in a Serpentinizing System"

**Supplementary Table Captions.**

**Supplementary Table 1. Concentrations of reactants and products for methane oxidation coupled to O2, NO3- and SO3-2. Estimated energy availability in unites kJ per mole electron and kJ per kg fluid.**

**Supplementary Table 2. Core methanotroph metabolism genes, annotations and BLASTp results.**

**Supplementary Table 3. Descriptive statistics and Mann-Whitney U test parameters to assess the significance of distribution differences of aerobic methanotroph phylotypes with respect to total ammonia nitrogen (TAN).**

**Supplementary Table 4. ASVs classified as methanotrophs, the taxonomic assignment and BLASTn results.**

**Supplementary Table 5. Relative abundance of ASVs classified as methanotrophs.**

**Supplementary Table 6. Relative abundance of methanotroph phylotypes by group.**

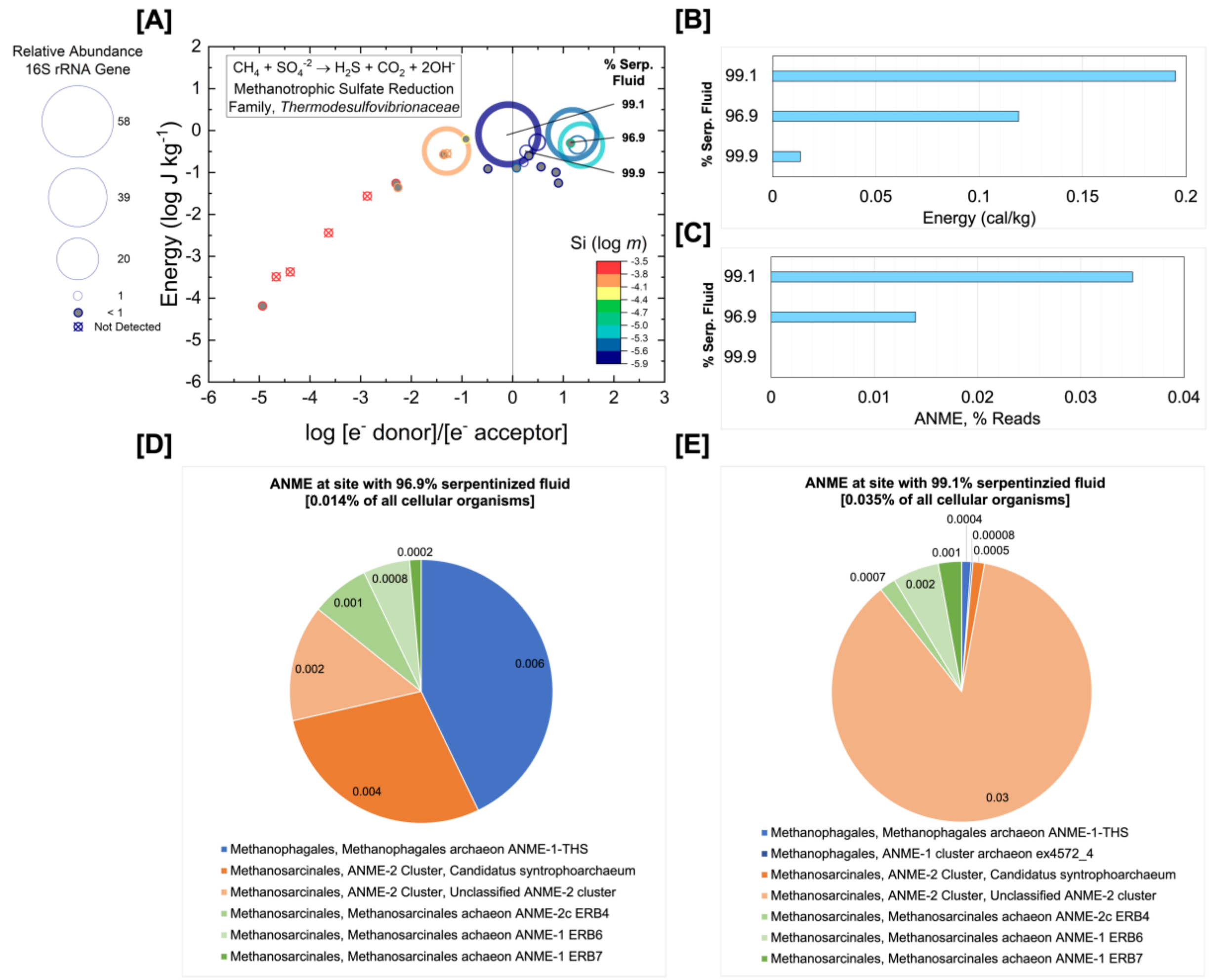


**Supplementary Figure 1. Potential for anaerobic methane oxidation coupled to sulfate reduction.**

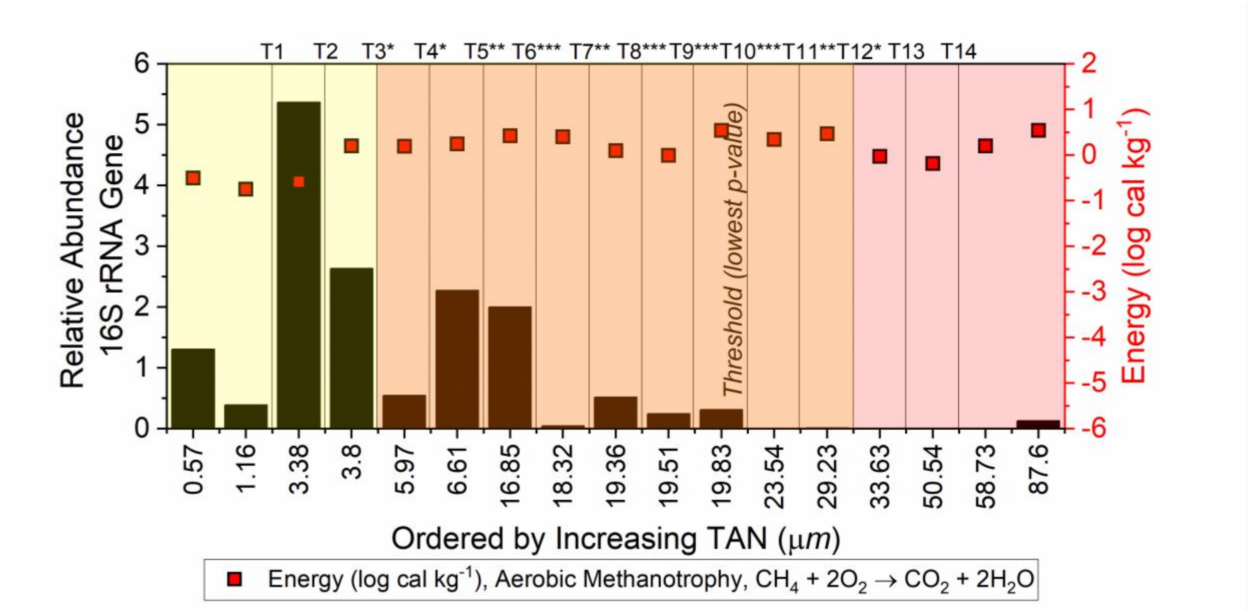


**Supplementary Figure 2. Visual summary of Mann-Whitney U-tests to assess distribution of aerobic methanotrophs with respect to total ammonia nitrogen (TAN).**

**Exsolution Calculation.**

To model the relative rates of $^{12}CH_4$ and $^{13}CH_4$ migration across the water-gas interface, we used the following equation,

$$\frac{dL}{dH} = \frac{1}{\alpha_k} \frac{(P\,\gamma - L_d)}{\left(P\,\gamma\,(\frac{H}{L})_g\,\alpha_{eq}\right) - \left(L_d\,(\frac{H}{L})_d\right)} \quad , \qquad (1)$$

where $\alpha_k$ is an experimentally determined kinetic isotopic fractionation factor, P is the average partial pressure of methane in the local atmospheric gas we measured, $\gamma$ is the Henry's law constant for methane that we obtained from SUPCRT92 (Johnson et al., 1992), $L_d$ is the concentration of the light isotope ($^{12}CH_4$) dissolved in solution, $(\frac{H}{L})_g$ is this average ratio of heavy to light methane in the local atmospheric gas, $(\frac{H}{L})_d$ is this ratio of heavy to light methane dissolved in solution, and $\alpha_{eq}$ is the equilibrium isotopic fractionation from dissolution. To constrain our upper bound for isotopic fractionation during exsolution we used an $\alpha_k$ value of 0.9990, representing no turbulence, and to constrain our lower bound we used a value of 0.9994, representing a high level of turbulence (Knox et al., 1992). Our values for $L_d$ and $(\frac{H}{L})_d$ for our upper bound were from sample 140115X while those values for our lower bound were from sample 140117F. Both sites that contained <1% shallow groundwater entrainment.
